# Over-Reliance on Prior Expectations in Relapsing-Remitting Multiple Sclerosis

**DOI:** 10.64898/2025.12.10.693435

**Authors:** Ahmad Pourmohammadi, Hesam Rezaei, Armin Adibi, Mina Dehghani, Arman Gorji, Sepehr Sima, Iman Adibi, Mehdi Sanayei

**Affiliations:** School of Cognitive Sciences, Institute for Research in Fundamental Sciences (IPM), Tehran, Iran; Department of Neurology, School of Medicine, Isfahan University of Medical Sciences, Isfahan, Iran; Center for Translational Neuroscience, Isfahan University of Medical Sciences, Isfahan, Iran; Isfahan Neuroscience Research Center, Isfahan University of Medical Sciences, Isfahan, Iran; School of Medicine, Isfahan University of Medical Sciences, Isfahan, Iran; Neuroscience and Neoplasia Artificial Intelligence Research Group (NAIRG), Hamadan University of Medical Sciences, Hamadan, Iran

**Keywords:** Multiple sclerosis, Bayesian models, cognitive impairments, aging

## Abstract

Cognitive impairment is a common and disabling feature of multiple sclerosis (MS). Bayesian models of perception and action provide a powerful framework to better understand how cognitive processes are altered in MS. In this case-control study, we employed a time reproduction paradigm within a Bayesian framework to investigate the underlying mechanisms of cognitive dysfunction in patients with relapsing-remitting MS. We applied a modified Bayesian observer model, which partitions time reproduction into three stages: sensory measurement, time estimation, and motor response. We found that both MS and control groups showed a systematic bias in reproducing time, overestimating short intervals and underestimating long intervals. This bias was significantly larger in patients with MS compared with controls, reflecting an over-reliance on prior expectations relative to sensory-motor information. Computational modeling indicated that this increased bias in the MS group was driven by greater measurement noise during the sensory stage. Moreover, central tendency bias increases with age in healthy participants as reliance on prior expectations becomes stronger than sensory-motor evidence. Interestingly, we found that this age-related effect on bias was absent in patients with MS. Further analysis showed both younger and older patients performed equally biased, and their performance was similar to older healthy participants.

## Introduction

Multiple sclerosis (MS) is a chronic inflammatory disease of the central nervous system and remains one of the leading causes of non-traumatic disability among young adults(Jakimovski et al., 2024). Disability in MS is not limited to physical symptoms. Cognitive impairment is prevalent in MS (Rao et al., 1991) and can lead to reduced quality of life, social functioning, and employment(A. Hakim et al., 2000). Previous studies have shown that cognitive impairment can begin early in the disease course, even before the onset of other clinical signs and symptoms(Cortese et al., 2016). Moreover, cognitive decline and disease progression can occur even in the absence of other neurological signs(Amato et al., 2001), underscoring the importance of cognitive assessment in patients with MS.

Despite the clinical importance of cognitive impairment in MS, its assessment and definition remain challenging. Systematic neuropsychological studies have demonstrated deficits across multiple domains, with impairments in information processing speed, memory, and executive functions consistently reported(Benedict et al., 2020; Rao et al., 1991). However, the binary classification of cognition as either “impaired” or “preserved” fails to capture the complexity of brain function and overlooks the dynamic nature of cognitive changes throughout the disease course. Moreover, MS is characterized by substantial heterogeneity in both clinical presentation and cognitive phenotypes(De Meo et al., 2021; Leavitt et al., 2018; Podda et al., 2021). Consequently, relying on a test targeting one specific domain may miss cognitive impairments in many patients. These limitations highlight the need to develop innovative approaches for cognitive evaluation in MS.

In this context, the Bayesian models of perception and action(Ma and Jazayeri, 2014) provide a promising approach to advance our understanding of cognitive impairments in MS. Although the external world is vast and full of information, our capacity to acquire sensory input is limited, and under many conditions our sensory measurements are noisy. To overcome these constraints, the brain generates prior expectations about the world and integrates them with the incoming sensory evidence to optimize perception and action. Over the past two decades, this framework has proven valuable for studying brain functions, supported by well-established behavioral paradigms and computational models.

In this study, we used the time reproduction paradigm within a Bayesian framework to investigate cognitive impairment in patients with MS. This approach is well suited to provide mechanistic insights into cognitive dysfunction in MS for several reasons. At the behavioral level, the task engages multiple processes, including sensory perception, motor reproduction, working memory and other cognitive functions(Jazayeri and Shadlen, 2010). Previous studies have shown this task engages multiple brain regions, reflecting the integration of distributed perceptual, cognitive, and motor systems(Jazayeri and Shadlen, 2015; Wang et al., 2018). At the computational level, well-established models allow task performance to be decomposed into distinct stages, enabling investigation of the mechanisms underlying different response patterns(Acerbi et al., 2012; Jazayeri and Shadlen, 2010).

We hypothesized that patients with MS would show impairments in the time reproduction task compared with healthy controls. Such deficits could arise from noisy sensory measurement, systematic over- or underestimation of time, or increased motor variability. To disentangle these possibilities, we applied a Bayesian observer model to identify the underlying mechanisms of cognitive impairments and to interpret them within the broader Bayesian framework. We found that patients with MS showed an over-reliance on prior expectations, driven by increased noise in the sensory measurement stage.

## Results

Participants completed a tablet-based time reproduction task in which they measured and reproduced time intervals between two visual flashes (Fig. 1). Five interval durations (0.4-1.9 sec) were presented pseudorandomly, and feedback was provided after each trial. For each participant, we performed a linear regression between presented and reproduced times, where the intercept captured overall over- or underestimation and the slope reflected sensitivity to duration changes and systematic bias. We then used a modified Bayesian observer model that divides time reproduction into three stages: sensory measurement, time estimation, and motor response. Each stage is described by one parameter: measurement noise (wₘ), gain factor (α), and motor noise (wᵣ), respectively.

**Figure 1.**
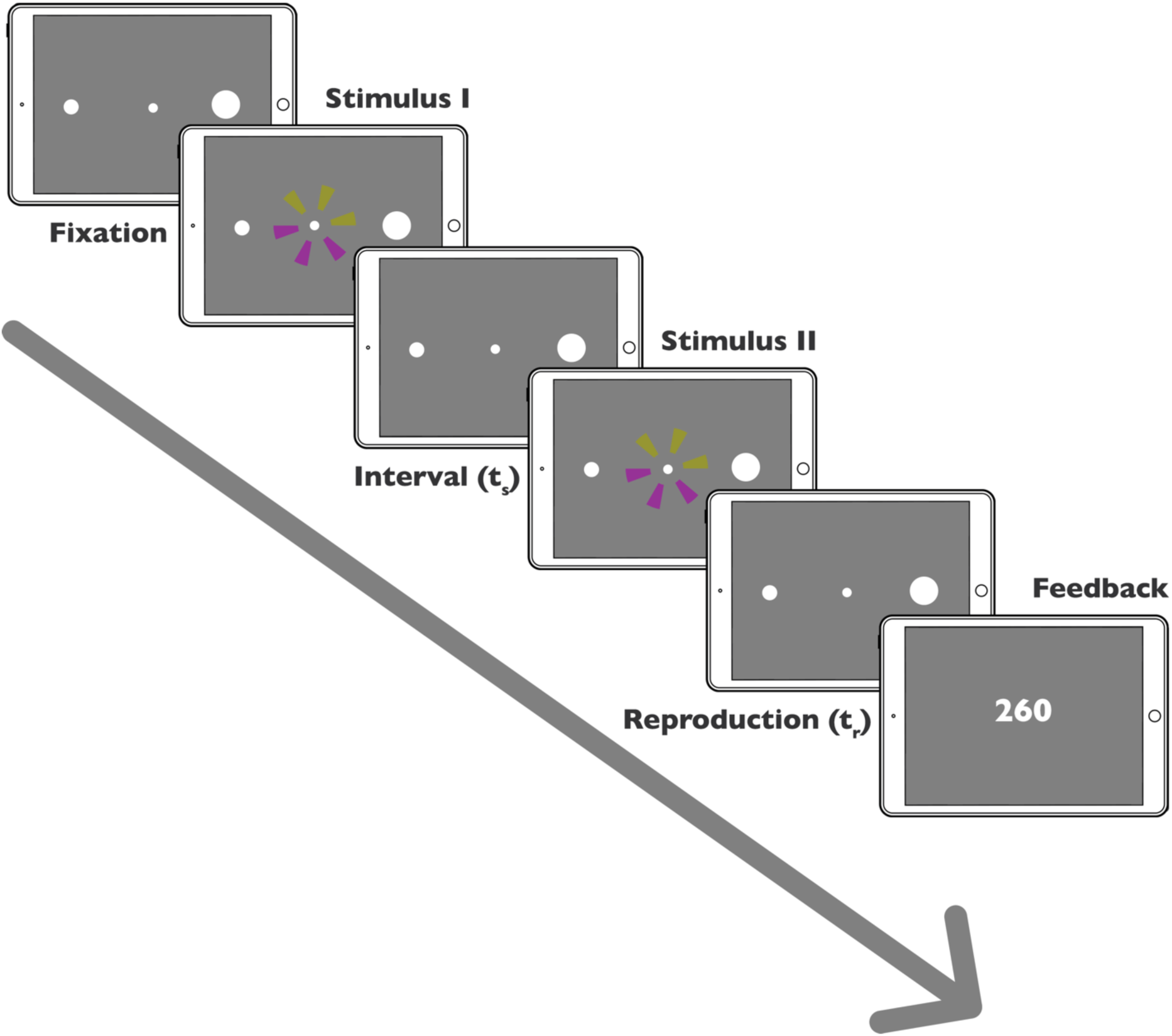
Time reproduction task paradigm. Each trial began with a fixation point and two peripheral targets (Fixation). After a random delay, the first wheel-like stimulus (Stimulus I) was briefly presented, followed by a sample interval (t_s_) and then a second stimulus (Stimulus II). Participants were instructed to measure the duration between the two flashes and reproduce it (t_r_) by tapping on the right side of the screen (Reproduction). At the end of each trial, feedback was provided showing the response error (t_r_ – t_s_) in milliseconds (Feedback).

We collected data from 92 relapsing-remitting MS patients and 149 control subjects. After exclusion, we enrolled 79 and 140 participants for the MS and control groups, respectively (See Methods for details). The MS and control groups did not differ significantly in age (MS: 36.33 ± 7.64 years; controls: 34.74 ± 8.12 years; p = 0.065), gender (MS: 73.42% female; control: 71.43% female; p = 0.87), or education (MS: 19.00% graduate degree; controls: 22.14% graduate degree; p = 1.00). Patients showed significantly slower performance on the 9-Hole Peg Test (9HPT; 24.72 ± 5.55 vs. 20.32 ± 2.93 sec, p < 0.0001) and reported higher anxiety levels (BAI; 12.76 ± 9.93 vs. 11.63 ± 9.50, p = 0.008). Other variables did not differ significantly between groups (Table 1).

**Table 1.** Demographic and clinical characteristics.

| Variable | MS (N=79) | Healthy (N=140) | p-value |
| --- | --- | --- | --- |
| Age | 36.33 ± 7.64 | 34.74 ± 8.12 | 0.065 |
| Graduate degree (%) | 15 (19.00) | 31 (22.14) | 1 |
| Female (%) | 58 (73.42) | 100 (71.43) | 0.87 |
| 9HPT | 24.72 ± 5.55 | 20.32 ± 2.93 | < **10<sup>-4</sup>\*** |
| HADS | 11.01 ± 7.67 | 10.84 ± 6.40 | 0.74 |
| MFIS_Physical | 12.57 ± 9.58 | 10.08 ± 7.40 | 0.10 |
| MFIS_Cognitive | 11.01 ± 9.06 | 9.31 ± 8.46 | 0.17 |
| MFIS_Psychosocial | 2.54 ± 2.54 | 3.08 ± 5.67 | 0.72 |
| Beck | 13.54 ± 10.35 | 11.63 ± 9.50 | 0.21 |
| BAI | 12.76 ± 9.93 | 9.21 ± 7.72 | **0.008\*** |
| FSS | 36.13 ± 15.68 | 34.67 ± 11.92 | 0.21 |
| EDSS | 1.06 ± 1.10 | - | - |

\* Denotes a significant difference (p-value < 0.05)

### Increased temporal bias in MS

We plotted the subjects’ average reproduced time against the presented time interval for the control (Fig. 2A) and MS (Fig. 2B) groups. We then applied a linear regression between the presented time and the reproduced time, separately for control and MS participants. In this task, the intercept of the regression reflects overall shifts in time reproduction: higher intercepts indicate a general tendency to overestimate, while lower intercepts suggest underestimation. The slope represents sensitivity to changes in duration; a slope of 1 reflects optimal timing, whereas deviations from 1 indicate systematic bias in reproducing shorter or longer intervals. The slope of the fitted line was smaller in the MS group (mean ± SD: 0.50 ± 0.26) than the control group (0.58 ± 0.22, t = 2.36, p = 0.027, Cohen’s d = 0.33; Fig. 2C) which means participants in MS group were less sensitive to time than control groups (or more biased towards the mean). The intercept of the fitted lines was not different between the two groups (MS = 0.51 ± 0.27, Control = 0.44 ± 0.23, U = 4716, p = 0.07; Fig. 2D).

**Figure 2.**
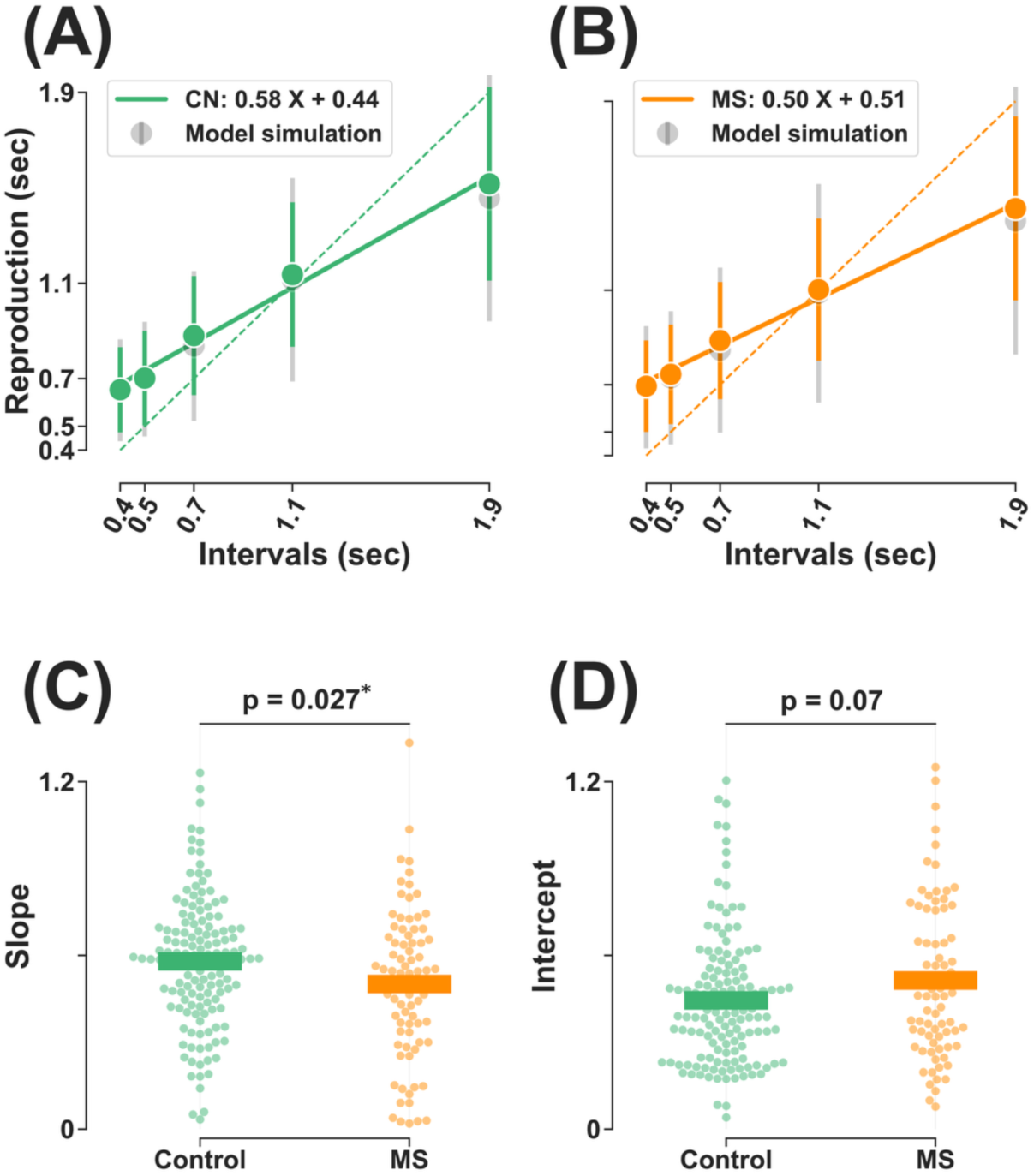
Time reproduction in MS and control groups. (A–B) Average reproduced time plotted against presented intervals for control (A, green) and MS (B, orange) groups. Solid circles indicate the mean reproduced time for each interval, with error bars showing the standard deviation. Solid lines represent the best-fit linear regressions for each group, with corresponding slope and intercept values shown in the legends. Gray circles and error bars represent the mean ± SD of model simulations. The dashed diagonal line indicates the identity line (x = y). Both groups exhibited systematic biases, with overestimation of shorter intervals and underestimation of longer intervals, as well as a scalar increase in variability with increasing interval duration. The Bayesian observer model captured both features in both groups. (C–D) Individual regression slopes (C) and intercepts (D) for control (green) and MS (orange) participants. The MS group showed significantly lower slopes compared to controls (p = 0.027), indicating reduced sensitivity (or bias towards the mean) to interval duration. Intercepts did not differ significantly between groups (p = 0.07). Horizontal bars denote group means.

### Increased measurement noise as a source of temporal bias in MS

To identify the source of decreased sensitivity (or increased bias towards the mean) in patients with MS, we applied a Bayesian observer model. This model allowed us to separately quantify variability at the time measurement stage (w_m_), estimation stage (α), and motor reproduction stage (w_r_). The model provided a good fit to the data for both the control (R² = 0.99) and MS (R² = 0.99) groups. We then compared the fitted parameters of the model between the two groups. The measurement noise (w_m_) was significantly larger in the MS group (0.38 ± 0.15) than in the control group (0.32 ± 0.11; U = 4328, p = 0.008, Cohen’s d = 0.44; Fig. 3A). In contrast, w_r_ did not differ between groups (Control: 0.24 ± 0.09; MS: 0.24 ± 0.10; U = 5713, p = 0.68; Fig. 3B). Similarly, α showed no significant difference between groups (Control: 0.98 ± 0.20; MS: 0.96 ± 0.26; U = 5414, p = 0.80; Fig. 3C).

**Figure 3.**
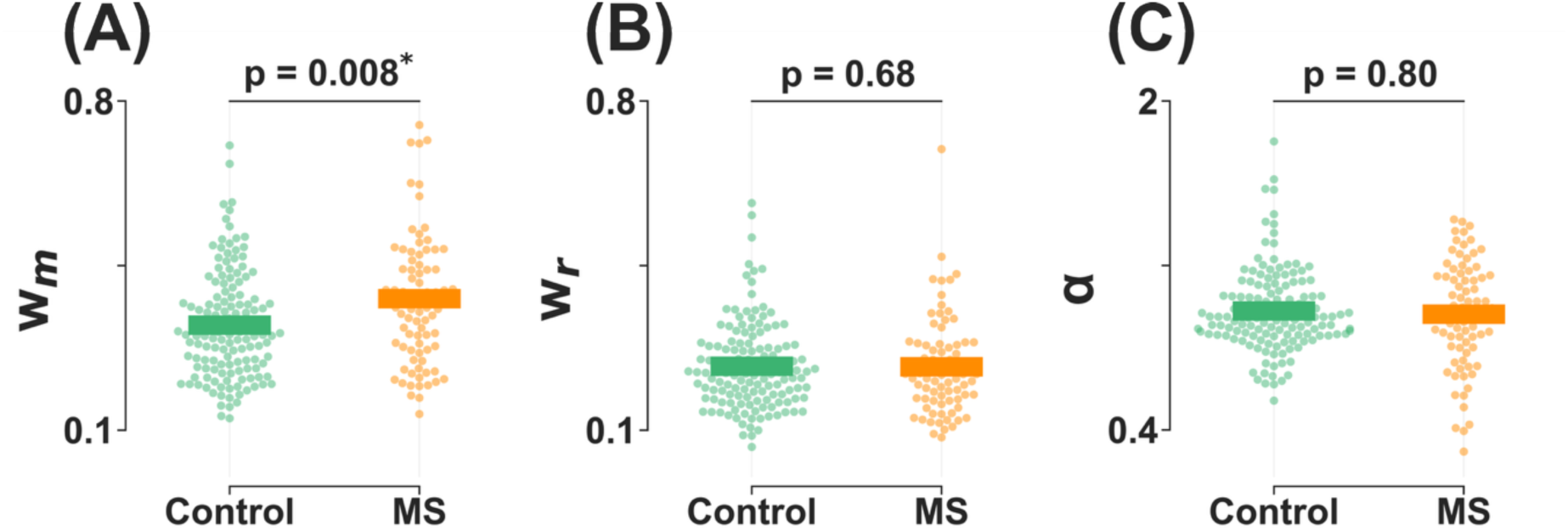
Bayesian observer model parameters in MS and control groups. (A–C) Parameters of the Bayesian observer model fitted to individual data: measurement noise (w_m_, A), motor noise (w_r_, B), and multiplicative gain factor (α, C). The MS group (orange) showed significantly larger w_m_ compared to controls (green, p = 0.008), whereas no group differences were observed for w_r_ (p = 0.68) or α (p = 0.80). Horizontal bars denote group means.

### Impact of demographic and clinical factors on time reproduction

We used univariate linear regression to assess the effects of demographic and clinical factors on two key outcomes that MS patients were worse than the control group, the slope of reproduction time and w_m_, within each group. Regression results are presented in Tables 2. None of the demographic or clinical variables were significantly associated with time reproduction slope or w_m_ in the MS group. In healthy participants, age was significantly associated with reproduction slope (β = −0.007, p-adj = 0.035) and w_m_ (β = 0.004, p-adj = 0.002). W_m_ was also significantly associated with education (β = −0.066, p-adj = 0.012) and gender (β = −0.063, p-adj = 0.012).

**Table 2.** Effects of demographic and clinical variables on time reproduction parameters.

|  | MS | Healthy |
| --- | --- | --- |
| Dependent | Independent variables | Independent variables |
| Slope | Age ($\beta = -0.008$, p-adj = 0.159) | **Age ($\beta = -0.007$, p-adj = 0.035\*)** |
| | Education ($\beta = 0.009$, p-adj = 0.948) | Education ($\beta = 0.091$, p-adj = 0.239) |
| | Gender ($\beta = 0.078$, p-adj = 0.647) | Gender ($\beta = 0.059$, p-adj = 0.346) |
| | 9HPT ($\beta = -0.011$, p-adj = 0.159) | 9HPT ($\beta = 0.007$, p-adj = 0.43) |
| | HADS ($\beta = 0.002$, p-adj = 0.888) | HADS ($\beta = -0.003$, p-adj = 0.43) |
| | MFIS_Physical ($\beta = 0.002$, p-adj = 0.888) | MFIS_Physical ($\beta = -0.005$, p-adj = 0.239) |
| | MFIS_Cognitive ($\beta = 0$, p-adj = 0.948) | MFIS_Cognitive ($\beta = -0.002$, p-adj = 0.43) |
| | MFIS_Psychosocial ($\beta = 0.002$, p-adj = 0.934) | MFIS_Psychosocial ($\beta = 0.002$, p-adj = 0.616) |
| | Beck ($\beta = 0.003$, p-adj = 0.554) | Beck ($\beta = -0.002$, p-adj = 0.43) |
| | BAI ($\beta = -0.003$, p-adj = 0.554) | BAI ($\beta = -0.002$, p-adj = 0.43) |
| | FSS ($\beta = -0.001$, p-adj = 0.799) | FSS ($\beta = -0.003$, p-adj = 0.239) |
| | EDSS ($\beta = -0.061$, p-adj = 0.159) | - |
| W<sub>m</sub> | Age ($\beta = 0.005$, p-adj = 0.134) | **Age ($\beta = 0.004$, p-adj = 0.002\*)** |
| | Education ($\beta = -0.065$, p-adj = 0.261) | **Education ($\beta = -0.066$, p-adj = 0.012\*)** |
| | Gender ($\beta = -0.064$, p-adj = 0.261) | **Gender ($\beta = -0.063$, p-adj = 0.012\*)** |
| | 9HPT ($\beta = 0.006$, p-adj = 0.157) | 9HPT ($\beta = -0.005$, p-adj = 0.287) |
| | HADS ($\beta = -0.0$, p-adj = 0.985) | HADS ($\beta = 0.001$, p-adj = 0.784) |
| | MFIS_Physical ($\beta = -0.0$, p-adj = 0.985) | MFIS_Physical ($\beta = 0.002$, p-adj = 0.252) |
| | MFIS_Cognitive ($\beta = -0.0$, p-adj = 0.985) | MFIS_Cognitive ($\beta = 0.0$, p-adj = 0.885) |
| | MFIS_Psychosocial ($\beta = -0.0$, p-adj = 0.985) | MFIS_Psychosocial ($\beta = -0.001$, p-adj = 0.784) |
| | Beck ($\beta = -0.001$, p-adj = 0.811) | Beck ($\beta = 0.0$, p-adj = 0.813) |
| | BAI ($\beta = 0.002$, p-adj = 0.577) | BAI ($\beta = 0.001$, p-adj = 0.784) |
| | FSS ($\beta = 0.002$, p-adj = 0.261) | FSS ($\beta = 0.001$, p-adj = 0.178) |
| | EDSS ($\beta = 0.038$, p-adj = 0.134) | - |

\* Denotes a significant difference (p-value < 0.05)

We then applied multivariate linear regression in the control group to assess the effects of age, gender, education, and hand motor performance (9HPT) on slope and w_m_. After adjustment for covariates, age remained the only significant predictor of both outcomes (Table 3).

**Table 3.** Effects of age on time reproduction parameters after controlling for sex, education, and hand motor performance.

| Dependent | Healthy |
| --- | --- |
|  | Multivariate regression model |
| Slope | Age ($\beta = -0.006$, p-adj = 0.029*) +<br>Education ($\beta = 0.051$, p-adj = 0.308) +<br>Gender ($\beta = 0.021$, p-adj = 0.649) +<br>9HPT ($\beta = 0.006$, p-adj = 0.399) |
| $W_m$ | Age ($\beta = 0.003$, p-adj = 0.007*) +<br>Education ($\beta = -0.034$, p-adj = 0.145) +<br>Gender ($\beta = -0.042$, p-adj = 0.055) +<br>9HPT ($\beta = -0.004$, p-adj = 0.145) |

\* Denotes a significant difference (p-value < 0.05)

To further investigate the effects of aging on time reproduction in MS versus control groups, we divided participants within each group into younger and older subgroups based on the median age of all participants (35 years; Fig. 4). A two-way ANOVA showed significant main effects of age (F = 6.50, p = 0.011) and group (F = 4.18, p = 0.042) on slope, with no significant interaction between age and group (F = 0.51, p = 0.47). Post-hoc pairwise comparisons showed the younger MS subgroup did not differ from the older control subgroup (t = 0.08, p-adj = 0.935; Fig. 4A), whereas the older MS subgroup showed significantly lower slopes than the younger control subgroup, with a larger effect size as expected (t = 3.03, p-adj = 0.020, Cohen’s d = 0.61; Fig. 4A). Similarly, a two-way ANOVA indicated significant main effects of age (F = 14.73, p < 0.001) and group (F = 6.97, p = 0.009) on w_m_, with no significant interaction (F = 0.06, p = 0.81). Post hoc comparisons followed the same pattern: the younger MS subgroup did not differ from the older control subgroup (U = 1074, p-adj = 0.89; Fig. 4B), while the older MS subgroup showed significantly higher w_m_ values than the younger control subgroup (U = 1029, p-adj = 0.001, Cohen’s d = 0.83; Fig. 4B).

**Figure 4.**
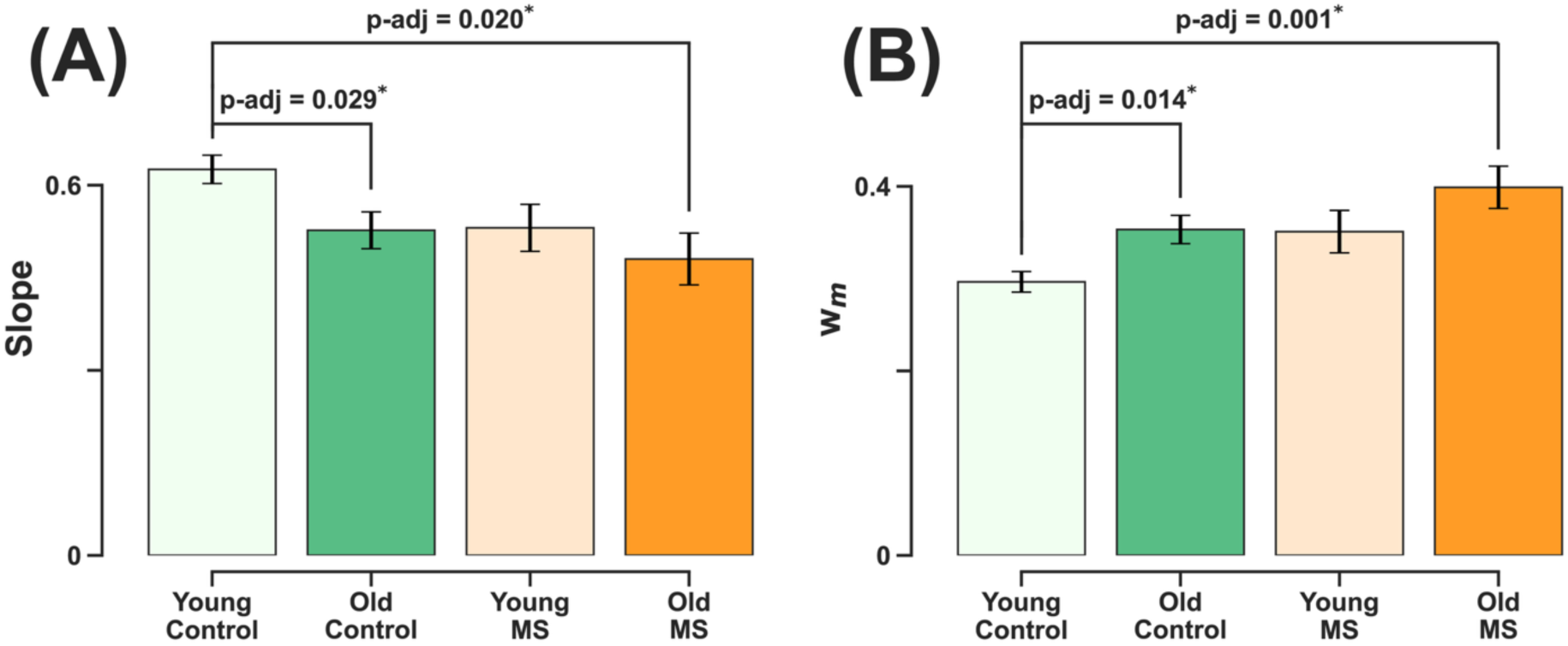
Effects of age on time reproduction in MS and control groups. Participants in each group were divided into younger and older subgroups based on the median age. (A) Slope was significantly larger in the young control subgroup compared to the old control (p-adj = 0.029) and old MS (p-adj = 0.020) subgroups. No significant differences were observed between young MS and old controls (p-adj = 0.93). (B) W_m_ was significantly larger in the young control subgroup compared to the old control (p-adj = 0.014) and old MS (p-adj = 0.001) subgroups. No significant differences were observed between young MS and old controls (p-adj = 0.89). Bars represent group means ± SEM.

## Discussion

We used the time reproduction paradigm within a Bayesian framework of perception to investigate cognitive impairment in patients with multiple sclerosis (MS). Both MS and control groups showed a systematic bias in reproducing time, overestimating short intervals and underestimating long intervals. This pattern, known as the central tendency bias, has been reported in time reproduction tasks and other perceptual domains(Lejeune and Wearden, 2009; Glasauer and Shi, 2021; Ma and Jazayeri, 2014). MS patients showed increased bias compared with controls. Using a modified Bayesian observer model, we demonstrated that this increased bias in the MS group was driven by greater measurement noise during the sensory measurement stage.

### Pathophysiological reliance on prior expectations

Based on Bayesian framework(Ma and Jazayeri, 2014; Shi et al., 2013), sensory processing involves combining information from the external world with prior expectations generated by the brain. Our results demonstrated an increased central tendency bias in patients with MS, indicating greater reliance on prior information than on sensory-motor information. This over-reliance may reflect the brain’s strategy under noisy conditions, where performance is optimized by assigning more weight to prior expectations relative to sensory-motor evidence. To place our findings in the larger context, it is helpful to consider evidence from other neurological and psychiatric conditions.

Previous studies using the time reproduction task in Parkinson’s disease have shown increased central tendency bias in patients(Bon-Mi Gu et al., 2015; Malapani et al., 2002, 1998; Terao et al., 2021). This increased bias has been reported during medication-off periods and improved after dopamine replacement therapy(Malapani et al., 1998) and deep brain stimulation of the subthalamic nucleus(Koch et al., 2004). An over-reliance on prior expectations has been also reported both in subclinical hallucination-prone individuals(Davies et al., 2018; Duhamel et al., 2023; Powers et al., 2017; Teufel et al., 2015) and in patients with schizophrenia experiencing hallucinations(Cassidy et al., 2018). Previous studies in Parkinson’s and psychotic disorders have proposed different non-mutually exclusive mechanisms to explain this over-reliance on prior expectations, which may also account for our findings in MS: (I) When sensory information is weak or noisy(Freedman et al., 1987; Javitt and Sweet, 2015), the brain compensates by relying more heavily on prior expectations. (II) Circuit-level changes may lead to an overweighting of prior information in perceptual inference, independent of sensory noise(Duhamel et al., 2023). (III) When motor responses are noisy or unreliable, the brain may rely more on prior expectations to stabilize motor output. (IV) Deficits in short-term or working memory may further increase dependence on prior expectations(Akrami et al., 2018; Dušek et al., 2012; Malapani et al., 2002; Terao et al., 2021).

In this study, we found that patients with MS showed increased measurement noise at the sensory stage, while motor noise remained comparable to that of healthy controls. The connection between our findings in MS and previous observations in neurodegenerative and psychotic disorders may reflect shared pathophysiological mechanisms underlying cognitive impairments across these conditions, highlighting an interaction between sensory processing, prior expectations, and working memory. Although we did not investigate the role of working memory in the present study, our group and others have consistently reported working memory dysfunction in MS(Motahharynia et al., 2023; Pourmohammadi et al., 2023), Parkinson’s disease(Ramos and Machado, 2021), and psychotic disorders(Forbes et al., 2009). Taken together, a potential explanation for the over-reliance on prior expectations in MS involves pathophysiological mechanisms that increase noise in sensory information, circuit-level changes that overweight prior information, and coexisting working memory dysfunction.

### Accelerated cognitive aging in MS and over-reliance on prior expectations

The connection between our results in MS and findings from other neurodegenerative disorders highlights a potential interaction between the over-reliance on prior expectations and neurodegenerative processes. Age is one of the key factors that influence neurodegeneration and disease progression in MS(Conway et al., 2019; Graves et al., 2023), and it provides an important context for interpreting our results. Our findings, along with previous studies (for review see(Turgeon et al., 2016)), showed that in healthy people, central tendency bias increases with age as reliance on prior expectations becomes stronger than sensory-motor evidence. Interestingly, we found that this age-related effect on bias was absent in patients with MS. Further analysis showed both younger and older patients performed equally biased, and their performance was similar to older healthy participants. Moreover, this effect is not limited to the time reproduction task. In previous studies, our team found that working memory precision decreases with age in both healthy individuals and patients with MS. We also showed that the source of working memory error in patients with MS is similar to that observed in older healthy adults(Esfahan et al., 2024; Motahharynia et al., 2023). These findings suggest an accelerated cognitive aging in patients with MS. Recent studies have also reported accelerated biological aging in patients with MS compared with healthy individuals(Siavoshi et al., 2025b, 2025a), and the accelerated cognitive aging observed in our study may represent one aspect of this broader aging process. Although our study included a limited sample size across different age ranges, it provides an initial step toward understanding the complex interaction between aging, MS, and cognitive decline.

In summary, we found that patients with MS showed an overreliance on prior expectations, driven by increased noise in sensory measurements. Moreover, MS appears to abolish the normal age-related increase in reliance on prior expectations, as both younger and older patients were equally biased, and their performance was comparable to that of older healthy participants.

## Materials and methods

### Study design and participants

We enrolled 92 patients with relapsing-remitting multiple sclerosis (MS), recruited through a full-census approach from the multiple sclerosis clinic at Kashani Hospital, Isfahan, Iran. The control group comprised 149 healthy individuals with no first-degree relatives diagnosed with MS.

Inclusion criteria consisted of a confirmed diagnosis of MS(Thompson et al., 2018), age between 18 and 55 years, an Expanded Disability Status Scale (EDSS) score between 0 and 6, and normal or corrected-to-normal visual acuity.

Exclusion criteria included a clinical relapse or corticosteroid use within three months prior to enrollment; major neurological disorders other than MS (e.g., stroke, epilepsy, brain tumors, or CNS infections); history of brain surgery; psychiatric illnesses (e.g., major depressive disorder, bipolar disorder, or schizophrenia); substance or alcohol abuse; acute febrile illness at the time of testing; chronic systemic conditions such as diabetes, renal or hepatic failure, chronic obstructive pulmonary disease (COPD), hypothyroidism, or hyperthyroidism; and abnormal performance on the Nine-Hole Peg Test (9-HPT > 45 sec).

The study was conducted in accordance with the Declaration of Helsinki and its subsequent revisions. Ethical approval was obtained from the Ethics Committee of Isfahan University of Medical Sciences (Ethics Code: IR.ARI.MUI.REC.1403.132).

### Time reproduction task

The time reproduction task was developed as and ipadOS app using Swift programming language and in Xcode (version 13) and was used on a 10-inch Apple iPad. Details of the task and its ipadOS application design have been described previously(Gorji and Fathi Jouzdani, 2024; Pourmohammadi and Sanayei, 2023).

The task paradigm is illustrated in Fig. 1. Each trial began with the presentation of a central fixation point (diameter: 0.2°, at 50 cm distance) and two peripheral targets (left target: 0.5° diameters, right target: 2° diameter, 10° eccentricity). The targets were not relevant to this task, though we kept them for consistency between this tasks and other timing tasks that are not reported here. After a random delay (0.5 sec plus a random time from 0 to 0.2 sec), two similar wheel-like stimuli (2.5° diameter) were flashed (for 33.3ms each) sequentially around the fixation point. The subject measured the time between two flashes, sample interval or t_s_, and produced a matching interval, t_r_, by tapping on the right side of screen. Across trials, t_s_ was sampled from one of 5 discrete values pseudorandomly (0.4, 0.5, 0.7, 1.1, 1.9 sec, uniform distributions). At the end of each trial, we showed the response error (t_r_ - t_s_) as feedback for 1 sec to the subject. The inter-trial interval was 2 sec, and each block contained 40 trials. Each subject participated in a short training and 6 blocks.

### Data collection and questionnaires

First, all participants received detailed instructions and provided written informed consent. They then participated in an interview to collect demographic and clinical information, followed by 9-HPT and EDSS administered by a neurologist. Subsequently, they completed a series of standardized self-report questionnaires in Persian, including Beck Depression Inventory-II (BDI-II)(Toosi et al., 2017), Beck Anxiety Inventory (BAI)(Kaviani and Mousavi, 2008), Hospital Anxiety and Depression Scale (HADS)(Montazeri et al., 2003), Modified Fatigue Impact Scale (MFIS)(Ghajarzadeh et al., 2013), and Fatigue Severity Scale (FSS)(Shahvarughi-Farahani et al., 2013).

### Statistical analysis

For each participant, we applied the 1.5 interquartile range (IQR) method to exclude outlier trials. We then performed a linear regression between the presented time and the reproduced time for each participant. In this task, the intercept of the regression reflects overall shifts in time reproduction: higher intercepts indicate a general tendency to overestimate, whereas lower intercepts suggest underestimation. The slope represents sensitivity to changes in duration (or bias towards the mean): a slope of 1 reflects optimal timing, while deviations from 1 indicate systematic bias in reproducing shorter and longer intervals.

A slope of zero or negative values indicates unreliable time reproduction for that participant. Based on this reliability criterion, we excluded 9 participants (out of 149, 6.67%, 2 males) from the control group, and 13 (out of 92, 16.46%, 1 male) form the MS group. The number of excluded participants was significantly higher in the MS group than in the control group (Chi-square, *χ* = 4.48, p < 0.05). The average percentage of excluded trials per subject, based on 1.5 IQR method, was 4.86 ± 2.23 % in the control group and 5.74 ± 3.00 % in the MS group which was not significantly different between the two groups (Mann-Whitney test, P = 0.06).

Statistical analyses were performed using Python 3.12.7, with statistical significance defined as p < 0.05. To compare the MS and control groups, we first assessed the normality of data distributions using the Shapiro-Wilk test. If the data were not normally distributed, we applied the Mann-Whitney U test; otherwise, we used the independent-samples t-test. For regression analyses, we used the *OLS* function from the *statsmodels* library in Python. Multiple comparisons were corrected using the false discovery rate (FDR) method.

### The Bayesian observer model

To investigate the sources of temporal error, we applied a modified version of the Bayesian observer model(Pourmohammadi and Sanayei, 2023; Jazayeri and Shadlen, 2010; Sima and Sanayei, 2024), which partitions time reproduction into three stages: sensory measurement, time estimation, and motor reproduction. In the measurement stage, the observer takes noisy measurements (tₘ) of the sample intervals (tₛ). Measurement noise (wₘ) is the model parameter corresponding to this stage.

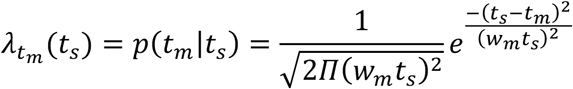

In the estimation stage, the Bayesian model combines the likelihood function with the prior and uses the mean of the resulting posterior distribution to generate an estimate (tₑ). In our model, Bayes least squares (BLS) was used as the mapping rule. The multiplication gain factor (α) in this stage represents systematic overestimation or underestimation.

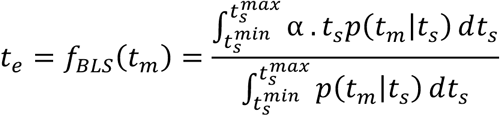

In the motor response stage, the observer uses t_e_ to reproduce noisy response, t_r_. Motor noise (w_r_) is the model parameter corresponding to this stage.

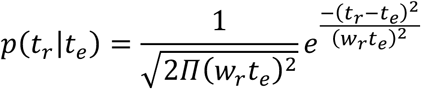

In our dataset, each trial comprised a sample time interval (tₛ) and the corresponding reproduced time (tᵣ). To implement the Bayesian observer model and capture the direct relationship between tₛ and tᵣ, we applied the following formulation.

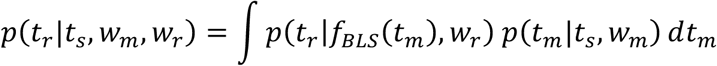

We used maximum likelihood estimation to fit the model and determine the optimal parameters (wₘ, wᵣ, α) for each participant. Model fitting was performed in Python using the SciPy library with the Nelder-Mead downhill simplex optimization method. To ensure robustness, the fitting procedure was repeated with multiple different initial parameter values.

### Language editing and artificial intelligence

A large language model (LLM) based tool (ChatGPT, OpenAI) was used to assist with grammar refinement and English language editing of the manuscript. Authors wrote the manuscript text and then used the following prompt for each paragraph:

“Any grammar or language mistakes and suggestions to improve language fluency of the text.”

All language suggestions were carefully reviewed, and the manuscript was revised independently by three authors to ensure accuracy and appropriateness.

## Data availability

The data that support the findings of this study are available from the corresponding author, upon request.

## Acknowledgements

None

## Supplementary material

None

